# A frameshift variant in the melanophilin gene is associated with loss of pigment from shed skin in ball pythons (*Python regius*)

**DOI:** 10.1101/2023.06.09.542917

**Authors:** Izabella Lederer, Basmah Shahid, Uyen M. Dao, Alex Brogdon, Hannah Byrtus, Marcelin Delva, Orson Deva, Paige Hatfield, Mikayla Hertz, Jenna Justice, Sarah Mavor, Erin Pilbeam, Zoe Rice, Abbey Simpson, Hallie Temar, Richard Wynn, Joana Xhangolli, Chiron W. Graves, Hannah S. Seidel

## Abstract

Melanophilin is a myosin adaptor required for transporting the pigment melanin within cells. Loss of melanophilin in fish, birds, and mammals causes pigmentation defects, but little is known about the role of melanophilin in non-avian reptiles. Here we show that a frameshift in the melanophilin gene in ball python (*P. regius*) is associated with loss of pigment from shed skin. This variant is predicted to remove the myosin-binding domain of melanophilin and thereby impair transport of melanin-containing organelles. Our study represents the first description of a melanophilin variant in a non-avian reptile and confirms the role of melanophilin across vertebrates.

## Introduction

The pigment melanin is responsible for brown and black skin colors throughout animals. Melanin in higher vertebrates is produced in cells called melanocytes, then transferred to cells called keratinocytes (Moreiras *et al*. 2021). Keratinocytes make up the bulk of the skin (Matsui and Amagai 2015), and melanin within these cells is essential for proper coloration of the animal (e.g. Ménasché *et al*. 2000; Matesic *et al*. 2001). Keratinocytes are continually sloughed off as dead skin, and melanin remains visible in the dead cells (Fig 1A).

**Figure 1.**
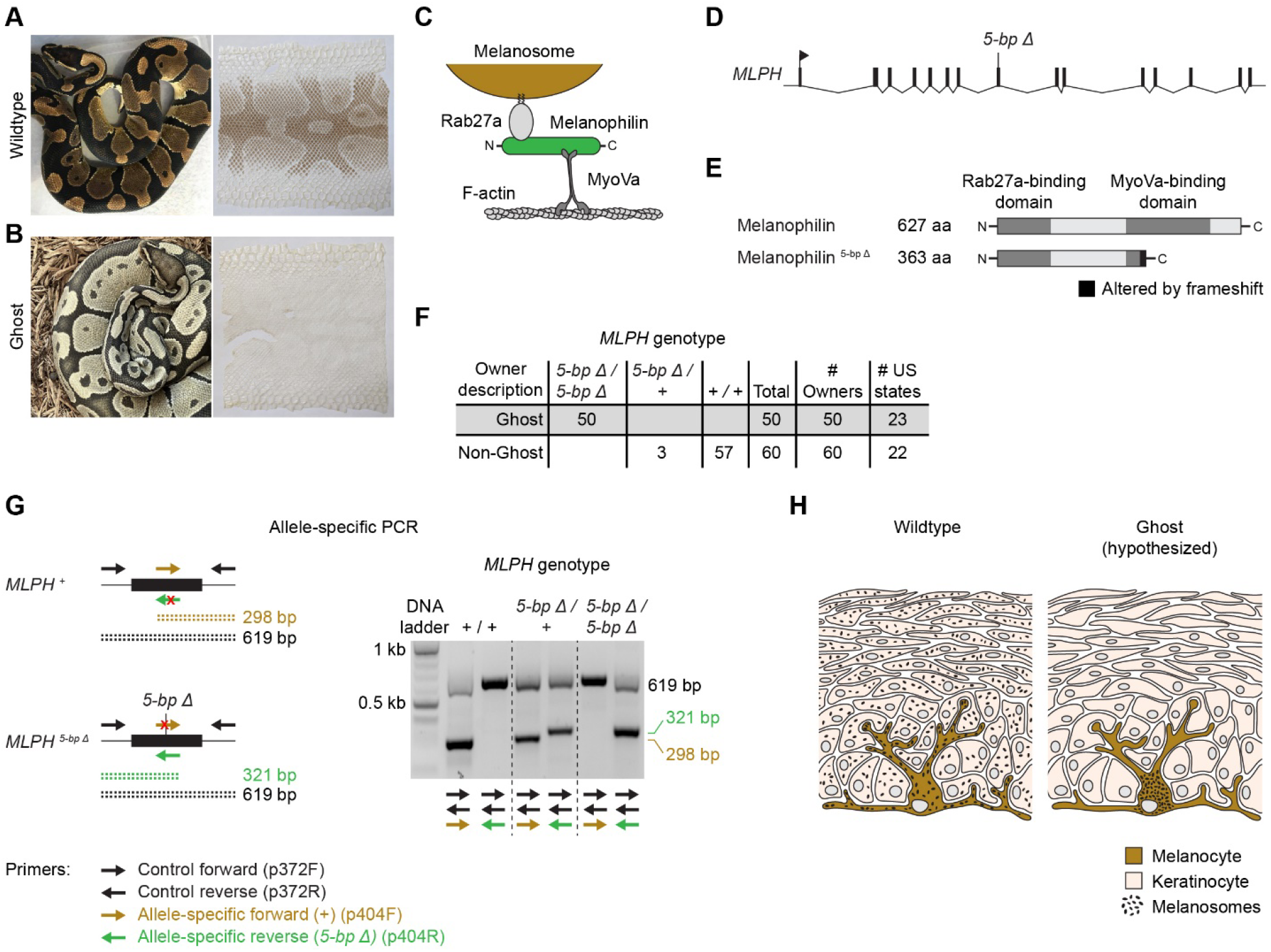
Loss of pigment in shed skin is associated with a 5-bp deletion in *MLPH*. (A-B) Wildtype and Ghost ball pythons and their shed skins. Photos, Chiron Graves and Thomi Gill. The wildtype photo is used with permission Brown et al. (2022). (C) Schematic of melanosome transport. (D) Structure of *MLPH*, including the 5-bp deletion found in Ghost animals. (E) Wildtype melanophilin protein (top) and melanophilin truncated by the 5-bp deletion (bottom). (F) Allele-specific PCR used to genotype the 5-bp deletion. Reactions used two control primers (black) and one primer specific for either the wildtype allele (brown) or the 5-bp deletion allele (green). (G) Association between the Ghost phenotype and the 5-bp deletion in *MLPH*. The number of US States indicates the total number of states from which sheds were collected. (H) Tissue architecture of the outer layer of the skin. In wildtype skin, melanosomes are present in melanocytes and keratinocytes. In Ghost skin, melanosomes are hypothesized to be absent from keratinocytes.

Transfer of melanin from melanocytes to keratinocytes occurs through transfer of melanin-containing organelles, called melanosomes (Le *et al*. 2021). Melanosomes are produced within the cell body of the melanocyte, which is located in the basal layer of the skin (Fig 1H). Mature melanosomes are trafficked from the cell body of the melanocyte to the tips of its dendritic processes, which extend to reach dozens of overlying keratinocytes (Hume and Seabra 2011; Fukuda 2021). Melanosomes are transferred to keratinocytes at these tips, through mechanisms that likely include exocytosis by the melanocyte and phagocytosis by the keratinocyte (Moreiras *et al*. 2021). Trafficking of melanosomes within the melanocyte is therefore essential for uptake of melanin by keratinocytes.

Trafficking of melanosomes within the melanocyte has been characterized in mammals, where it requires three proteins: MyoVa, Rab27a, and melanophilin (Figure 1C). MyoVa is a motor protein that transports melanosomes along actin filaments (Mercer *et al*. 1991). Rab27a is a small GTP-binding protein anchored in the membrane of the melanosome (Wilson *et al*. 2000; Hume *et al*. 2001; Wu *et al*. 2002b). Melanophilin is a linker protein that attaches Rab27a to the myosin motor (Hume *et al*. 2002; Wu *et al*. 2002a; Fukuda *et al*. 2002; Nagashima *et al*. 2002; Strom *et al*. 2002). Melanophilin is thought to be dedicated to trafficking of melanosomes, whereas MyoVa and Rab27a play broader roles in trafficking additional types of vesicles in other cell types (Langford 2002; Fukuda 2013)

Consistent with the role of MyoVa, Rab27a, and melanophilin in melanosome trafficking, loss of any of these proteins causes defects in pigmentation. Loss of melanophilin in zebrafish prevents melanosomes from dispersing within melanocytes, thereby impairing adaptation to low light conditions (Sheets *et al*. 2007). Loss of melanophilin in mammals and birds impairs transfer of melanin to keratinocytes, resulting in lighter skin, hair, and feathers (e.g. Vaez *et al*. 2008; Manakhov *et al*. 2019). This loss contributes to breed-specific coloration in a number of domesticated species, including chicken (Vaez *et al*. 2008), quail (Bed’hom *et al*. 2012), cat (Ishida *et al*. 2006), dog (Drögemüller *et al*. 2007; Bauer *et al*. 2018; Van Buren *et al*. 2020), rabbit (Lehner *et al*. 2013; Fontanesi *et al*. 2014; Demars *et al*. 2018), mink (Cirera *et al*. 2013; Manakhov *et al*. 2019), and cattle (Li *et al*. 2016). Similar defects in pigmentation upon loss of MyoVa or Rab27a, but with additional defects in neurons and immune cells (Pastural *et al*. 1997; Ménasché *et al*. 2000). Little is known about melanosome trafficking outside mammals, birds, and zebrafish, including in non-avian reptiles.

An emerging model for genetic studies in reptiles is the ball python (*P. regius*) (Brown *et al*. 2022; Garcia-Elfring *et al*. 2023; Dao *et al*. 2023). Ball pythons are native to west Africa, but have become common as pets. The pet population includes many heritable variants of the normal color pattern. One variant, known as Ghost, shows a phenotype suggesting a defect in melanosome trafficking. Ghost animals have lighter brown-to-black coloration compared to wildtype animals, and their shed skin is largely devoid of pigment (Fig 1B). Shed skin is primarily composed of keratinocytes, and absence of pigment in these cells suggests a defect in the transfer of melanin to keratinocytes. Ghost animals are healthy and show no abnormalities outside pigmentation. The Ghost phenotype is recessive, and the causative variant is unknown. (Alternate names for the Ghost phenotype are Hypo and Orange Ghost. For simplicity, we use the term Ghost to refer to any of these terms.)

The goal of the current study was to identify the genetic cause of the Ghost phenotype. We hypothesized that this phenotype was caused by a loss-of-function variant in a gene controlling melanosome trafficking. Obvious candidates were the melanophilin gene, the MyoVa gene, or the Rab27a gene. We prioritized the melanophilin gene because Ghost animals are healthy and lack the severe neurological or immunological impairments characteristic of loss of MyoVa or Rab27a (Pastural *et al*. 1997; Ménasché *et al*. 2000). Our approach was to collect samples of pet ball pythons (shed skins) from across the United States and search for putative loss-of-function variants in the melanophilin gene. A similar approach has been used previously by our group to identify variants controlling melanin production (Brown *et al*. 2022) and patterning (Dao *et al*. 2023).

## Results

We hypothesized that the Ghost phenotype was caused by a loss-of-function variant in the melanophilin gene (*MLPH*). To search for putative loss-of-function variants in *MLPH*, we amplified and sequenced the coding regions and adjacent splice sites of *MLPH* in one animal described by its owner as Ghost and one animal having normal coloration (henceforth ‘wildtype’). We then compared sequences from the two animals. We observed that the Ghost animal was homozygous for a 5-bp deletion in the eighth coding region of *MLPH* (OR035642:c.4919_4923del, Fig 1D). No other coding variants and no splice-site variants were observed. The 5-bp deletion creates a frameshift in the transcript (p.Gln350Ilefs*15), which is predicted to truncate the protein at 363 amino acids, compared to 596 amino acids in wildtype (Fig 1E). This truncation removes most of the MyoVa binding domain, which is required for melanosome trafficking (Hume *et al*. 2006). We conclude that the 5-bp deletion likely disrupts protein function and might be causative for the Ghost phenotype.

To test whether the 5-bp deletion in *MLPH* was associated with the Ghost phenotype in a broader population of animals, we genotyped 110 additional animals for the deletion. These animals consisted of 50 animals described by their owners as Ghost and 60 animals as described as Non-Ghost. The Non-Ghost animals included wildtype animals, as well as animals exhibiting color variants other than Ghost. Animals within each phenotypic category were collected from unique owners and therefore represent a broad sampling of the pet population. We observed that all animals described as Ghost were homozygous for the 5-bp deletion (Fig 1G). Non-Ghost animals, by contrast, did not carry the deletion (57 animals) or were heterozygous for it (three animals) (Fig 1G). These data show that homozygosity for the 5-bp deletion in *MLPH* is perfectly associated with the Ghost phenotype.

## Discussion

We propose that the 5-bp deletion in *MLPH* is causative for the loss of pigment from shed skin observed in Ghost animals. The 5-bp deletion is predicted to produce a truncated melanophilin protein that lacks most of the MyoVa-binding domain (Fig 1E). The 5-bp deletion might also cause the melanophilin transcript to be degraded by nonsense-mediate decay. Truncation of melanophilin at a similar position in mink is associated with lighter fur color (Cirera *et al*. 2013; Manakhov *et al*. 2019), consistent with truncations at this position disrupting melanosome trafficking. Likewise, removal of the MyoVa-binding domain in mammalian cells causes melanosomes to remain clustered within the cell body of the melanocyte (Hume *et al*. 2006), where they cannot be transferred to keratinocytes (Wu *et al*. 2012). We predict that a similar outcome occurs in Ghost animals. We propose that melanosomes in Ghost animals remain clustered within the cell body of the melanocyte and are never transferred to keratinocytes (Fig 1H). This hypothesis provides a simple explanation for the absence of pigment in shed skin in Ghost animals.

A limitation of our study is that we only examined a single candidate gene—the melanophilin gene. We cannot formally exclude the possibility that variants in other regions of the genome contribute to the Ghost phenotype. We view this possibility as unlikely, however, given that the Ghost phenotype exactly matches the predicted loss-of-function phenotype of the melanophilin gene.

The melanophilin gene has been repeatedly targeted by artificial selection in domesticated birds and mammals to produce breed-specific coloration (e.g. Ishida *et al*. 2006; Vaez *et al*. 2008; Bed’hom *et al*. 2012; Manakhov *et al*. 2019). This gene represents a good target for artificial selection because loss of the melanophilin protein alters pigmentation without affecting the overall health of the animal. Our study suggests that artificial selection for loss of melanophilin extends from birds and mammals to pet snakes.

## Methods

### Shed collection

Ball python sheds were recruited from pet owners and breeders by placing announcements social media. Contributors were instructed to allow sheds to dry (if wet) and to ship sheds via standard first-class mail. To maximize genetic diversity within our sample, we included a maximum of one Ghost animal and one Non-Ghost animal per contributor. The Ghost category included animals described by their owners as Ghost (24 animals), Hypo (18 animals), or Orange Ghost (nine animals). The Non-Ghost category included animals described by their owners as wildtype (14 animals) or as having color phenotypes other than Ghost (47 animals). Sheds were inspected to confirm that sheds in the Ghost category were devoid of pigment and that sheds in the Non-Ghost category contained pigment. One shed was excluded from the study because it did not meet this criterion.

### Performing experiments in an undergraduate laboratory course

This study was performed as part of an undergraduate laboratory course at Eastern Michigan University. To avoid student errors in a classroom setting, the following precautions were implemented. First, students never handled more than one shed skin at the same time. Second, students performed negative and positive control reactions for all PCR amplifications. Data from students having incorrect controls were excluded from analysis. Third, all sequence reads were examined independently by three or more students. If the results of all students did not all agree, sequences were examined by the instructor (HSS).

### Gene annotation for *MLPH*

We based our gene annotation of *MLPH* in ball python on the genome of Burmese python (*Python_molurus_bivittatus-5*.*0*.*2*), the closest relative for which genome sequence was available (Castoe *et al*. 2013). (We did not attempt to recover *MLPH* transcripts in ball python because our samples were limited to shed skins, which do not contain intact mRNA.) The Burmese python genome is a scaffold-level assembly, and its gene annotations are imperfect (Brown *et al*. 2022). We therefore manually curated a gene annotation for *MLPH. MLPH* in Burmese python is annotated as having multiple isoforms, the longest of which (isoform X1) has 17 coding regions (NCBI *Python bivittatus* Annotation Release 102, 2018-05-24). We examined these coding regions and concluded that 15 of them were likely annotation correctly and two were likely intronic regions that had been misannotated as coding. This conclusion was based on the following evidence: (i) 15 of the 17 coding regions are well supported by RNA-seq exon coverage available in NCBI and encode protein sequences that are highly conserved with MLPH protein sequences from other vertebrates, including mouse (NP_443748.2), chicken (NP_001108552.1), anole lizard (XP_008112807.1), corn snake (XP_034282850.1), and western garter snake (XP_032069882.1), (ii) the two remaining coding regions, which represent coding regions 13 and 14 of *MLPH* isoform X1 (XM_025171758.1 / XP_025027526.1), are not well supported by RNA-seq coverage, and their protein sequences are absent from melanophilin proteins in other vertebrates, (iii) the region homologous to coding region 13 is fixed for a stop codon in ball python (n = seven animals, including five wildtype animals), and(iv)coding region 14 is unusually small (12 nucleotides), below the size needed to support robust splicing (Ustianenko *et al*. 2017). Our gene annotation for *MLPH* in ball python consisted of the 15 coding regions conserved across vertebrates. The exon-intron structure this annotation matches *MLPH* isoform X4 in Burmese python (XM_025171774.1 / XP_025027542.1), *MLPH* in corn snake (XM_034426959.1 / XP_034282850.1), and *MLPH* in western garter snake (XM_032213991.1 / XP_032069882.1).

### Primers for sequencing *MLPH* coding regions

Primers for sequencing *MLPH* coding regions were designed against the genome of Burmese python. These primers were the following: coding region 1, p383F (5’-AAT CAT TCA CGG CAG GCA AA-3’) and p383R (5’-TGG TGG TGG CAT GGA ATA CT -3’); coding region 2, p398F (5’-TCA CCT CCT CTC ACA GTC CA-3’) and p398R (5’-AAG GCT ATG GTT TGC AGT GC-3’); coding region 3, p385F (5’-TCC ATG TCT GAA GGG TCA CC-3’) and p385R (5’-TGA AGC ACC CAA GAC AGT CA-3’); coding region 4, p386F (5’-GCC TAA AGA TGT GCT GCT CC-3’) and p386R (5’-AGG CCA TCA AAA TCA GAC GC-3’); coding region 5, p399F (5’-GGA TGG CAG CAA TTG AAC CT-3’) and p399R (5’-ACC TGG CCG ATC TTA TCT GG -3’); coding region 6, p388F (5’-TGT GCT GTA TTC TGG CCT GA-3’) and p388R (5’-GCC CAC TCA CTG ACC TAT GT-3’); coding region 7, p389F (5’-CCA GGT CTA TGC CCA GTG AA-3’) and p389R (5’-GTT GTG ACG TGA AGA GCT GC -3’); coding region 8, p372F (5’-CCT GAA TCT CAA AGG GGT TAC A -3’) and p372R (5’-CAA TTT GTT GTG GGC TTA GTG -3’); coding region 9, p390F (5’-ATG CAG CCT TCC TCC TTG AT-3’) and p390R (5’-CTT GGG AGC AGT ACT GAG GA-3’); coding region 10, p400F (5’-AGT TCA AAA CAG TCA TGG GGA-3’) and p400R (5’-TAT GCA CAC TGG GGA TGA GG-3’); coding region 11, p401F (5’-TCC CTA CTG CTG AAC TGA CA-3’) and p401R (5’- TGT TTG TTG TGA ATC GCC CA-3’); coding region 12, p393F (5’-TTC CAC CAC TAG CAC CTG AC-3’) and p393R (5’-GAG CCT GGA GTG TAT CTT CCT-3’); coding region 13, p394F (5’-GCC TGT TTT GAC ACT TTT GCA-3’) and p394R (5’-GTC TTC AAG GCC CTT TCA GC-3’); coding region 14, p396F (5’-ACC GGT TCT TTC CCC ATT CT-3’) and p396R (5’- TGC TCC TTC CTC CTT GTG TT-3’); and coding region 15, p402F (5’-CGG GGA AAA TGG GTG AAC AA-3’) and p402R (5’-ACT GTG CAT AAA TCT CCA GGG A-3’). Amplicons were sequencing using p372F, p383R, p385R, p386R, p388F, p389F, p390R, p393F, p394R, p396F, p398R, p399F, p400F, p401F, and p402F.

### DNA extraction

Sheds containing visible dirt or debris were rinsed in tap water until clean. Sheds were air dried and lysed overnight at ∼60°C in ∼1 ml lysis buffer (100 mM Tris-HCl pH 8.0, 100 mM EDTA, 2% sodium dodecyl sulfate, 3 mM CaCl_2_, 2 mg/ml Proteinase K) per palm-sized piece of shed. Lysate was separated from visible fragments of undigested shed and further cleared by centrifugation at 13,000 x g for 2 min. To precipitate protein, ammonium acetate was added to supernatant to a final concentration of 1.875 M. Samples were incubated on ice for 5 min and centrifuged at 13,000 x g for 3 min at 4°C. Supernatant was mixed with an equal volume of magnetic bead mixture (10 mM Tris-HCl pH 8.0, 1 mM EDTA, 1.6 M NaCl, 0.2% Tween-20, 11% polyethylene glycol, 0.04% washed SpeedBeads [Sigma #GE45152105050250]) and shaken for 5 min. Beads were separated from supernatant using a magnet, washed twice in 0.2 ml 70% ethanol for 2 min, and air dried for ∼1 min. DNA was eluted from beads in TE buffer (10 mM Tris-HCl pH 8.0, 1 mM EDTA) at 65°C for >5 min.

### PCR and Sanger sequencing

PCR was performed using OneTaq polymerase (NEB #M0480). Reactions consisted of 1X OneTaq Standard Reaction Buffer, 200 μM dNTPs, 0.2 μM of each primer, and 0.025 U/μl OneTaq polymerase. Thermocycling conditions were as follows: 94°C for 2 min; 30 cycles of 94°C for 30 sec, 56-57°C for 30 sec, and 68°C for 1-2 min; and 68°C for 5 min. Reactions used 10-100 ng template DNA per 20 μl volume.

PCR products were purified for Sanger sequencing using magnetic beads. PCR reactions were mixed with three volumes of magnetic-bead mixture (10 mM Tris-HCl pH 8.0, 1 mM EDTA, 1.6-2.5 M NaCl, 0.2% Tween-20, 11-20% polyethylene glycol, 0.04% washed SpeedBeads [Sigma #GE45152105050250]) and agitated for 5 min. Beads were separated from supernatant using a magnet, washed twice in 0.2 ml 80% ethanol for >2 min, and air-dried for 1 min. PCR products were eluted from beads in 10 mM Tris-HCl pH 8.0 for >3 min at 65°C. Sanger sequencing was performed by Eton Bioscience Inc (etonbio.com).

### Protein domains

Locations of the Rab27a-binding domain and MyoVa-binding domain were determined by aligning the melanophilin protein sequence from ball python to the melanophilin protein sequence from mouse (NP_443748.2). This alignment was created using Clustal Omega (Sievers *et al*. 2011). In mouse, the Rab27a-binding domain corresponds to residues 1-150, and the MyoVa-binding domain corresponds to residues 300-484 (Hume *et al*. 2006). In ball python, the Rab27a-binding domain corresponds to residues 1-150, and the MyoVa-binding domain corresponds to residues 313-521.

### Genotyping the 5-bp deletion in *MLPH*

Animals were genotyped for the 5-bp deletion in *MLPH* using a pair of allele-specific PCR reactions (Fig 1F). Each PCR reaction contained three primers: two control primers (p372F, 5’-CCT GAA TCT CAA AGG GGT TAC A -3’; p372R, 5’-CAA TTT GTT GTG GGC TTA GTG -3’) and one primer specific for either the wildtype allele (p404F, 5’-GCG GGC GGG CAG GAG CCA AGC ATC ATC-3’) or the 5-bp deletion allele (p404R, 5’-GCG GGC GGG TTT TCT GTG ATG ATG CTC CT-3’). Both reactions produced a 619-bp product regardless of genotype. An additional, allele-specific product was produced from p404F for the wildtype allele (298 bp) and from p404R for the deletion allele (321 bp). Both reactions used an annealing temperature of 56°C and an extension time of 1 min.

## Acknowledgments

We thank the Educational Course Support program of New England BioLabs for reagents used in undergraduate teaching labs. Shed skins were contributed by the following individuals: Abby Kruse, Adam and Nicole Schmid, Adam Shirah of Proper Royals, Adrian Hoverter, Alycia Butler, Amanda Hall, Andelyn Czajka, Andy Rose, Bankrupt Reptiles, Big D, Bryan Rivera, Chelsey Wheeler, Chiron Graves, Chris Shelly, Chun Ku, CPJ Reptiles, Curtis Morgan, Cynthia Jones, Dale Porcher, Dave Dunn of Ball in Hand Pythons, David Burstein, Denise Jones, Duane McFalcand, Epic Vibrant Balls, Eric Chung, Erin Burt, Everything’s Better Orange, FNA Balls, Jaden Christensen, Jake and Ashley Brewster, Jake Lewis, Jamie Palazzo, Jason Roberts, Jeff Kearns, Jeff Linton, Jeromy Shaffer, JMB Exotics, Joe and Wendy Ducos, Joe Myers, John White, Justin Orbach of Just Incrediballs, Kai Li, Kandie Tucker, Kelly West, Kelsi Greene, Korie Dy, Kyle Reinke, Laniel Wolff, Leah Alexander, Lindsey VanOrman, Lynnet Melton, Manuel San Juan, Maryann Barbon, Matilda (Tillie) Groff, Matt Burton, Matt Huck, Michael Cole, Michael Nino, Miranda Joy Turpin, Morgan Evans, Morph Mechanics, Nate Fitzpatrick of Baltimore Balls, Nathan Granoff, Nick Deinhart, Ozzy Maldonado, Pacific Rim Serpents, Pedro Alequin of Imperial Morphs, Peter Holden, Phil Snyder, Randy Remington, Redstrom Reptiles, Reggie’s Urban Jungle, Roaring Pythons, Royal Black Balls, Ryan Boyd and Brittney Delacruz, Sean Strait of The Morph Lab, Sergio McDole, Shemier Bernard, Tammy Hutchinson, Taylor and Amanda Yereance, Tia Vogel,, Tyler Wilkins, Wes Mosher, Zac Parpart, and several anonymous contributors.

## Data availability

All relevant data are within the paper. DNA sequences are available from GenBank, under accession number OR035642.

